# Leg stiffness adjustment during hopping by dynamic interaction between the muscle and tendon of the triceps surae

**DOI:** 10.1101/2024.04.24.589455

**Authors:** Kazuki Kuriyama, Daisuke Takeshita

## Abstract

The biomechanics underlying bouncing exercises are characterized by the spring-like behavior of the human leg. However, the mechanism underlying the mechanistic contribution of muscle dynamics to the adjustment of leg stiffness is unclear. This study aimed to elucidate the mechanisms governing the changes in leg stiffness during hopping at different frequencies by examining the dynamics of the muscle–tendon complex (MTC) of the medial gastrocnemius muscle (MG). We hypothesized that an increase in muscle stiffness would augment leg stiffness, thereby enabling hopping at higher frequencies. Kinematic and kinetic data were obtained using a motion capture system and force plates. Simultaneously, ultrasound images of the MG were acquired to quantify the muscle fascicle length and pennation angle. The results showed that the stiffness of the MTC increased with hop frequency and exhibited a strong correlation with the leg stiffness. In addition, with increasing frequency, the fascicle contractions shifted from isometric to concentric. To explain these results, an MTC model comprising a contractile component (CC) and series elastic component (SEC) was constructed. We observed a negative CC stiffness, which increased the MTC stiffness. Although this result appears to diverge from our initial hypothesis, the effect of negative CC stiffness on MTC stiffness can be understood, from the perspective of two springs in series, as an extension of the very high stiffness effect. This quantitative understanding of the dynamic interaction between the muscle and tendon offers a unified framework for interpreting various results of previous studies on fascicle dynamics during hopping.

## 1. Introduction

The spring-like behavior of the human leg is a fundamental aspect of the biomechanics involved in movements with a stretch-shortening cycle (SSC), such as running and hopping. The dynamics of a bouncing body are often conceptualized through the widely employed “spring-mass model,” which models the body as a mass and the leg as a supporting spring (Blickhan, 1989). Using this model, the adaptability of leg spring stiffness, commonly referred to as “leg stiffness,” has been evaluated. Various studies revealed that leg stiffness can be adjusted in response to diverse conditions such as different ground surfaces (Ferris et al., 1998; Ferris and Farley, 1997), running speeds (Arampatzis et al., 1999; Garciá-Pinillos et al., 2019; Morin et al., 2006), and hop heights and frequencies (Farley et al., 1991; Farley and Morgenroth, 1999). Therefore, the ability to regulate leg stiffness is crucial to ensure stable movement in various environments.

Considerable research has been conducted on leg stiffness during hopping, one of the simplest SSC movements, suggesting the importance of adjusting leg stiffness to control hop frequencies (Brughelli and Cronin, 2008; Butler et al., 2003; Struzik et al., 2021). The adjustment of leg stiffness during hopping is primarily achieved by adjusting ankle stiffness rather than knee or hip stiffness (Farley and Morgenroth, 1999; Ferris et al., 1998; Hobara et al., 2011; Kuitunen et al., 2011). This implies that the triceps surae muscle–tendon complex (MTC) is involved in regulating leg stiffness, because ankle stiffness primarily depends on the stiffness of the triceps surae MTC, which comprises the largest plantar flexor muscle (Sakanaka et al., 2018). Moreover, MTC stiffness can be posited to be governed by the dynamics of muscle fibers (Monte et al., 2021; Van Hooren and Bosch, 2016). Using ultrasound techniques, previous studies reported variations in the dynamics of the triceps surae muscle during hopping at different hop frequencies and heights, where MTC stiffness is expected to differ (Hoffrén et al., 2012, 2015; Jessup et al., 2023; Sano et al., 2013). Therefore, the dynamics of triceps surae muscle fibers play an important role in regulating MTC stiffness and thereby leg stiffness. However, the mechanistic contribution of muscle dynamics to the adjustment of leg stiffness remains poorly understood.

In this study, we explain the adjustment of MTC stiffness, using an approach based on the classical MTC model comprising a contractile component (CC) and series elastic component (SEC) connected in series (Hill, 1938; Wakeling et al., 2023). Assuming that both the CC and SEC function as linear springs, the overall MTC stiffness (*k*_*MTC*_) can be expressed as

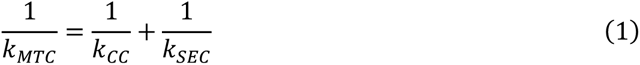

where *k*_*CC*_ and *k*_*SEC*_ represents the CC and SEC stiffnesses, respectively. MTC stiffness primarily depends on the CC stiffness when the CC is compliant, and it approaches the SEC stiffness when the CC becomes stiff (Fukashiro et al., 2001; Shorten, 1987). In other words, the CC must be stiff when a high MTC stiffness is required.

This study aimed to explain the changes in leg stiffness during hopping at different hop frequencies by examining the dynamics of the MTC of the medial gastrocnemius (MG) muscle. We anticipated that at lower hop frequencies, the muscle would behave as a compliant spring, elongating owing to the applied load. Conversely, the muscle was anticipated to behave similarly to a stiff spring, resist load, and remain isometric at higher hop frequencies. Therefore, we hypothesized that an increase in CC stiffness would augment leg stiffness and result in faster movements at higher hop frequencies.

## 2. Methods

### 2.1. Participants

Nine physically active male participants (age: 21.9±2.1 years; height: 170.4±4.4 cm; body mass: 59.0±3.6 kg) were enrolled in the study. Informed consent was obtained from each participant prior to their participation. This study adhered to the Declaration of Helsinki for studies involving human participants and was approved by the Ethics Committee of the University of Tokyo.

### 2.2. Task and procedure

After warm-up exercises, the participants performed repetitive hopping on force plates (Force Plate 9281E; 9281B; Kistler, Switzerland) using both feet at seven different frequencies ranging from 2.0 to 3.5 Hz, with increments of 0.25 Hz. These frequencies were determined using a metronome sound, and the measurements were performed in ascending order starting from the lowest frequency. This order was selected based on preliminary experiments, which indicated the need to ease the difficulty of trials involving finely distinguished frequencies. Once the exercise reached a steady state, data were collected for approximately 10 s. The participants were instructed to minimize the upper limb effect by placing their hands on opposite shoulders, maintaining their knees as extended as possible, and minimizing the duration of ground contact.

A three-dimensional motion capture system (MAC3D System, Motion Analysis, USA) comprising 10 infrared cameras was used to capture the coordinates of anatomical landmarks throughout the body at 200 Hz. Simultaneously, ground reaction forces (GRFs) were recorded at 2000 Hz using force plates and synchronized with the kinematic data. A real-time B-mode ultrasound apparatus (ARIETTA 70; FUJIFILM Healthcare, Japan) was used to capture longitudinal images of the right medial gastrocnemius muscle (MG) at 80 Hz. The ultrasound probe was securely fixed to the MG muscle belly using a styrofoam fixture to minimize movements. Synchronization of the ultrasound images with other data was achieved through an external trigger signal. An overview of the experimental setup is shown in Fig. 1.

**Fig. 1.**
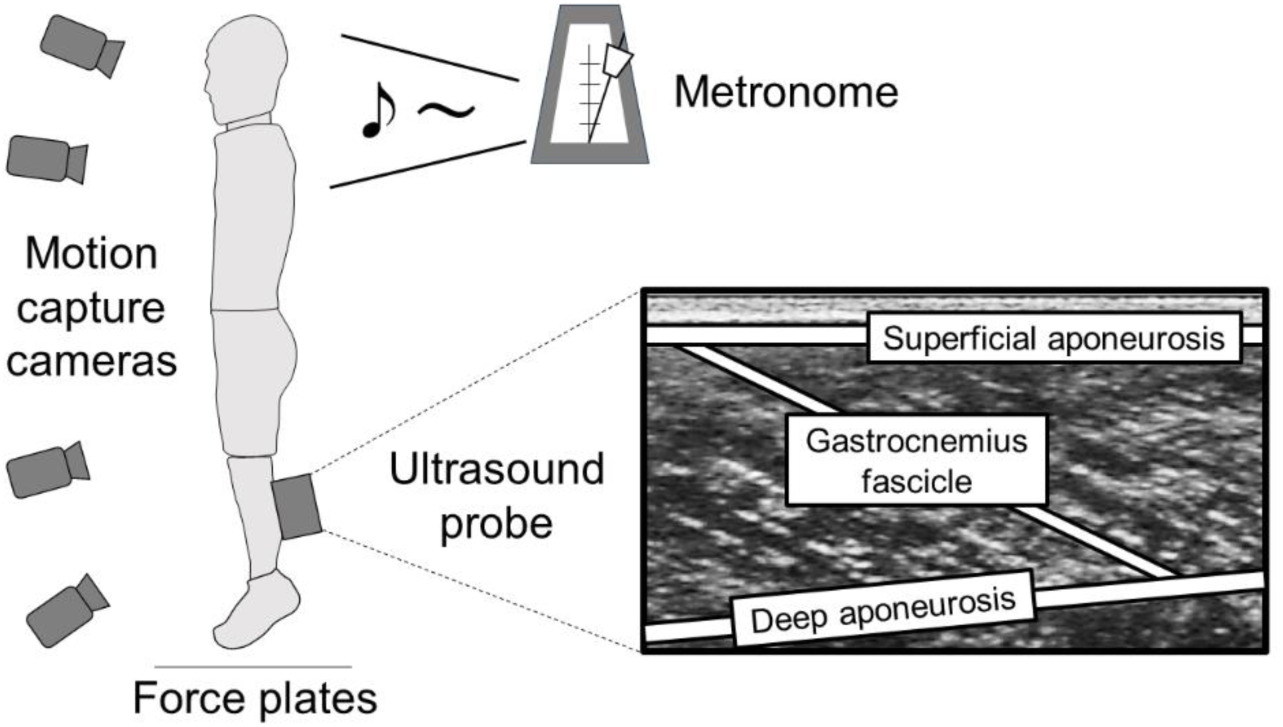
Overview of the experimental setup.

### 2.3. Data analysis

The dataset was subsequently analyzed using MATLAB 2023a (MathWorks, USA). For each hop frequency, the average of 10 hops was used for further analysis. Each hop was time-normalized using linear interpolation, with 0% and 100% representing the ground and subsequent ground contacts, respectively. The ground contact and takeoff timings were determined using GRFs with a threshold set at three times the standard deviation during the aerial phase (approximately 10 N).

Kinematic data were subjected to a fourth-order zero-lag low-pass Butterworth filter with a cutoff frequency of 10 Hz, which was approximately the average cutoff frequency determined through residual analyses (Winter, 2009) for each dimension and marker. Joint angle data were obtained from sagittal plane kinematic data by averaging the left and right marker coordinates. Joint torques were calculated using inverse dynamics (Winter, 2009), considering the inertial parameters of a Japanese body (Ae et al., 1992). The center of mass (CoM) of the entire body was computed from the CoM of each body segment.

The MTC length of the MG muscle (*L*_*MTC*_), representing the length from the origin to the insertion of the muscle, was determined using the following equation proposed by Hawkins and Hull (1990):

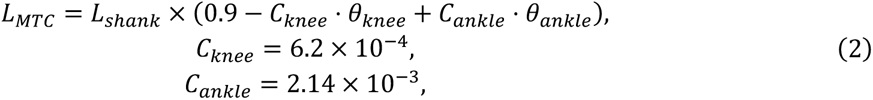

where *L*_*shank*_, *θ*_*knee*_ and *θ*_*ankle*_ represent the shank length, knee angle, and ankle angle, respectively, of each participant. Furthermore, the moment arm of the Achilles tendon was determined as *d*_*ankle*_ = *L*_*shank*_ ⋅ *C*_*ankle*_ ⋅ 180/*π*, based on the relationship between the ankle angle displacement and MTC length change (Bobbert et al., 1986).

The superficial and deep aponeuroses and MG fascicles were digitized from ultrasound images. The fascicle length was defined as the length along the fascicles from the superficial to deep aponeuroses, and the pennation angle was defined as the angle between the fascicles and deep aponeuroses (Van Hooren et al., 2020). These data were also filtered using a fourth-order zero-lag low-pass Butterworth filter with a cutoff frequency of 10 Hz, consistently with the kinematic data.

### 2.4. Statistics

The statistical significance was set at *p* < 0.05 . All data are presented as means ± standard deviations.

### 2.5. Calculation of stiffness

The leg stiffness was defined as the peak vertical GRF divided by the vertical displacement of the CoM between the ground contact and the moment of the peak GRF (McMahon and Cheng, 1990). Similarly to the leg stiffness, the CC, SEC, and MTC stiffnesses were defined as the peak load divided by the change in length between the ground contact and the moment of the peak load. The load on the MTC (*F*_*MTC*_ ) was calculated by dividing the ankle torque (*M*_*ankle*_) by the moment arm of the MTC (Kubo et al., 1999). The CC and SEC lengths (*L*_*CC*_ and *L*_*SEC*_, respectively) were calculated as the change in length along the longitudinal axis based on the fascicle length *L*_*fascicle*_ and pennation angle *⍺* (Fukunaga et al., 2001; Ishikawa et al., 2003):

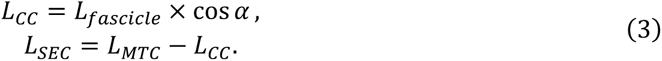

The loads on the muscles and tendons were assumed to be the same as those on the MTC. Fig. 2 shows a visual representation of each variable.

**Fig. 2.**
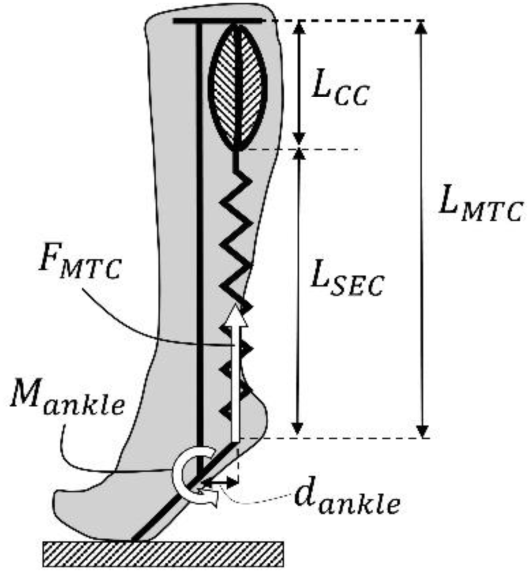
Model of the triceps surae MTC. *L*_*CC*_ , *L*_*SEC*_, and *L*_*MTC*_ represent the length of the contractile component (CC), series elastic component (SEC), and MTC, respectively; *d*_*ankle*_ represents the moment arm of the Achilles tendon; *M*_*ankle*_ represents the ankle joint torque; and *F*_*MTC*_ represents the load on the MTC.

Note that according to our definition, the CC stiffness may be negative, although the stiffness of a spring is typically positive. This peculiar behavior arises from the active force exerted by the CC, which can shorten against the increasing load.

## 3. Results

### 3.1. Hop frequencies

The determinant coefficients, calculated using the instructed and observed hop frequencies as the independent and dependent variables, respectively, ranged from 0.964 to 0.999 for each participant, indicating that the participants closely adhered to the instructed hop frequencies.

### 3.2. Correlation between leg and MTC stiffnesses

The leg and MTC stiffnesses increased with increasing hop frequency for most participants (Fig. 3). In addition, a significant positive correlation was observed between the two stiffness values for each participant, and the average correlation coefficient across the participants was 0.94.

**Fig. 3.**
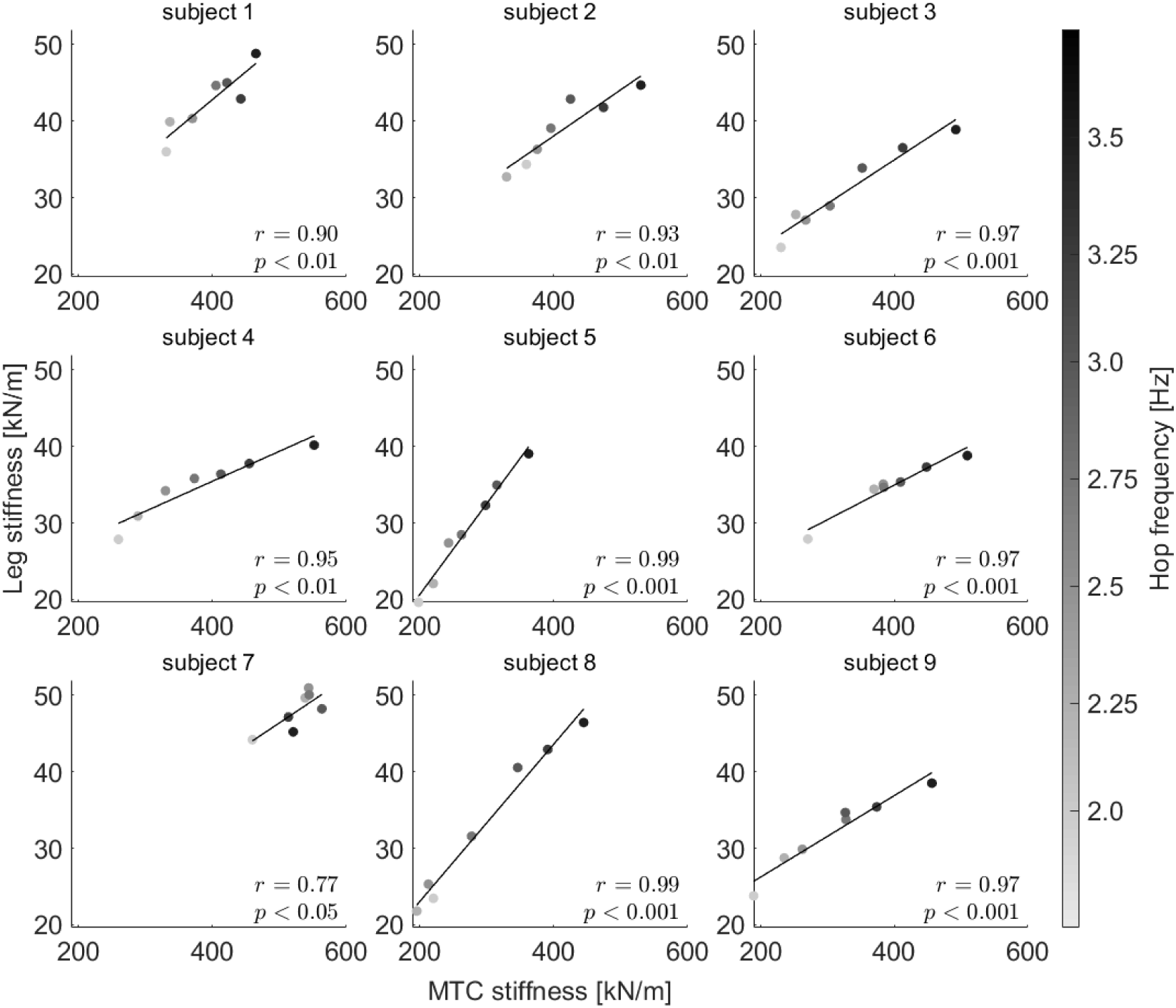
Correlation between MTC and leg stiffnesses for each participant. The darker and brighter dots correspond to higher and lower hop frequencies, respectively.

### 3.3. Contact and aerial durations

The contact duration remained relatively constant across the hop frequencies, whereas the aerial duration decreased with increasing hop frequency (Fig. 4). One-way analysis of variance (ANOVA) revealed no statistically significant main effect of hop frequency on contact duration (*p* = 0.68). Fig. 4 also shows the contact duration of the spring-mass model, in which the mass and stiffness are the average body mass and leg stiffness at 2 Hz, respectively. In this model, the contact duration increases with the hop frequency until motion becomes impossible to generate.

**Fig. 4.**
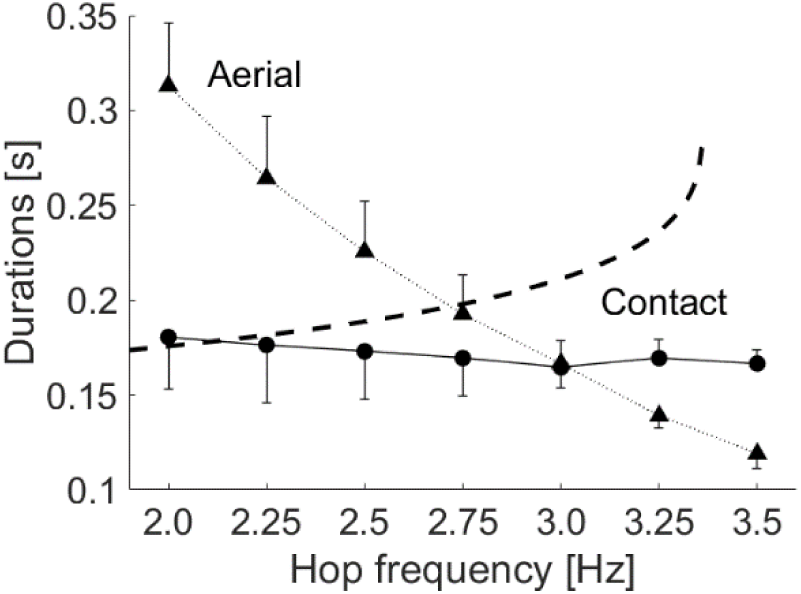
Ground contact (square) and aerial (triangle) durations at each hop frequency. The dashed line represents the contact durations for the spring-mass model, with constant stiffness value at 2.0 Hz. The data are shown as means and standard deviations.

### 3.4. Length changes of the MTC and fascicle

During ground contact, the MTC length exhibited consistent elongation followed by shortening, regardless of the hop frequency, whereas the fascicle contractions shifted from isometric to concentric with increasing hop frequency (Fig. 5(A)). The change in the MTC length was larger at lower frequencies and smaller at higher frequencies. The higher the frequency, the longer the fascicle length at the moment of ground contact and the greater the amount of shortening. Consequently, the ratio of the change in the fascicle length to that in the MTC length during ground contact was larger at higher frequencies (Fig. 5(B)).

**Fig. 5.**
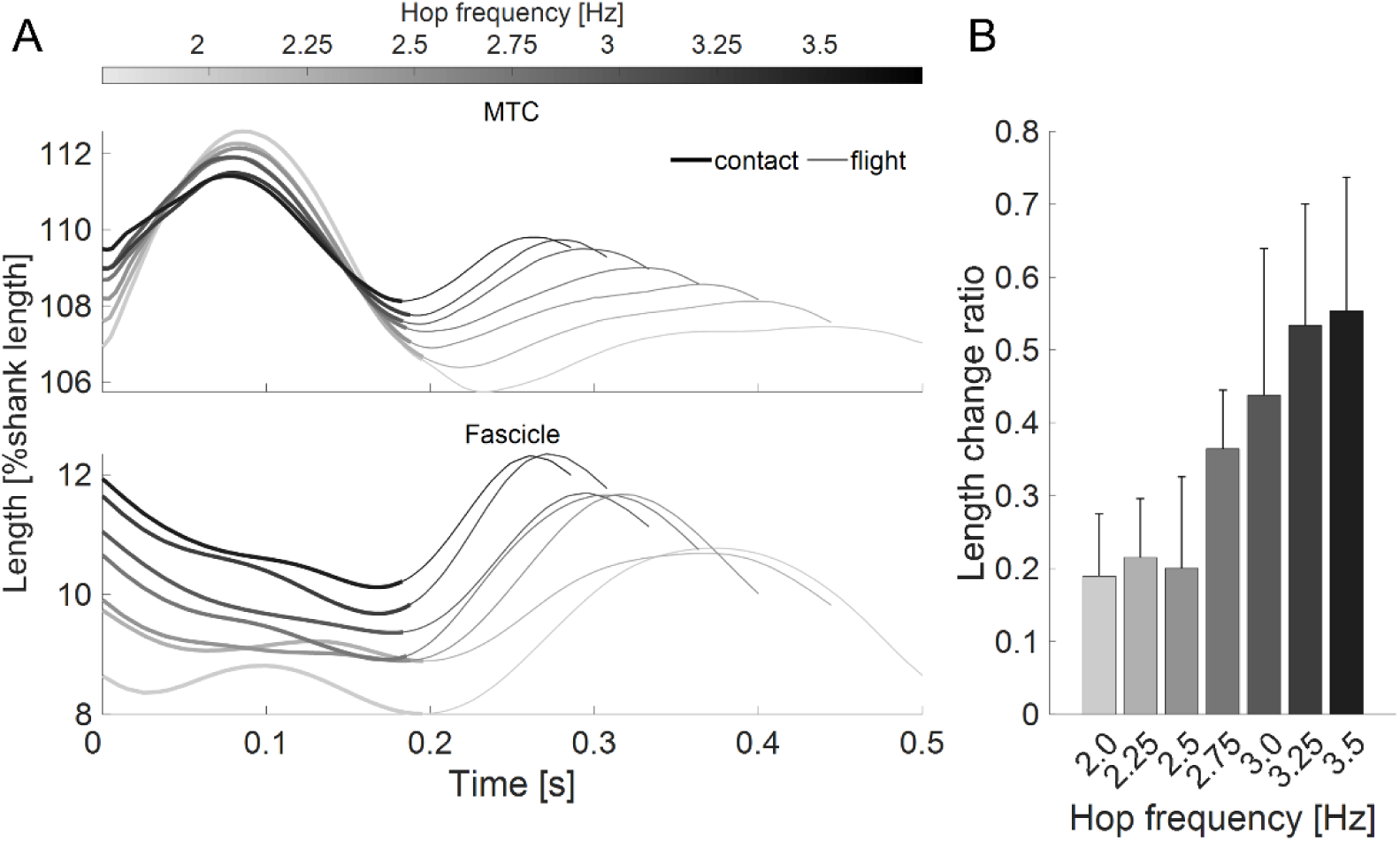
Average time course of the MTC and fascicle length during one cycle of hopping at different hop frequencies (A) and the ratio of changes in the fascicle length during ground contact relative to that of the MTC (B). Each length is shown relatively to the shank length. The darker and brighter lines correspond to higher and lower hop frequencies, respectively. The 0 second represents the ground contact, and the ends of the bold and thin lines represent the takeoff and subsequent ground contact, respectively.

### 3.5. Inverse of MTC, CC and SEC stiffnesses

Under most hop frequency conditions, the fascicle shortened during the first half of the grounding phase (Fig. 5(A)), resulting in a negative CC stiffness. The inverse of CC stiffness exhibited a larger negative value with increasing hop frequency (Fig. 6). The SEC stiffness remained almost constant, regardless of the hop frequency. One-way ANOVA showed a main effect of hop frequency on the inverse stiffness of the MTC and CC (*p* < 0.001) but no significant effect on that of the SEC (*p* = 0.74). The average SEC stiffness across the participants was 288±48 kN/m, ranging from 218 to 376 kN/m.

**Fig. 6.**
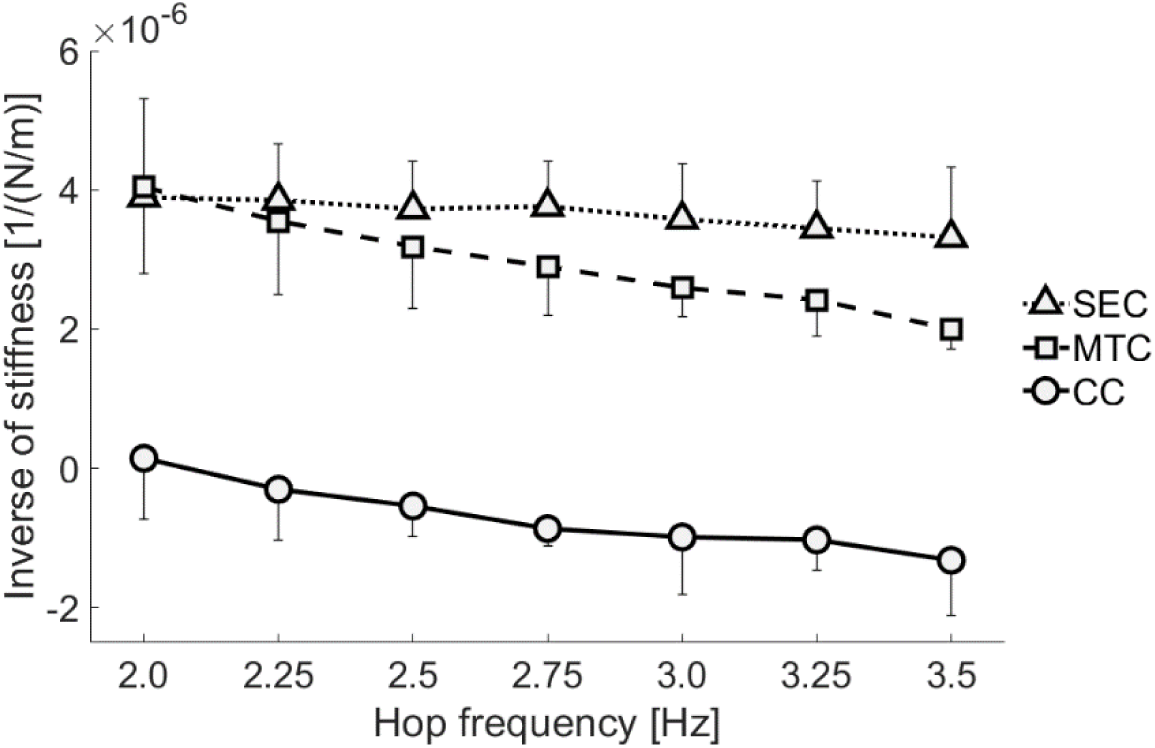
Inverse of stiffness values (see Eq. 1) at different hop frequencies. The square, triangle, and circle represent the stiffness values of MTC, SEC, and CC, respectively. The data are shown as means and standard deviations.

## 4. Discussion

This study was aimed at elucidating the mechanism underlying the changes in leg stiffness during hopping by examining the dynamics of the plantar flexor muscle. The major findings of this study can be summarized as follows: (1) The leg stiffness was highly correlated with the MTC stiffness, and (2) the MTC stiffness increased with hop frequency, primarily because of the negative CC stiffness. These results suggest that the effect of muscle dynamics on the MTC stiffness during SSC exercises can be quantitatively explained by considering the CC and SEC as two springs connected in series.

The results showed a consistent SEC stiffness, independent of hop frequency; this suggests the validity of our MTC model comprising the CC and SEC and assuming that the CC reflects muscle fascicle dynamics whereas the SEC functions as a purely passive component. Therefore, the consistent stiffness value of the SEC indicates the robustness of the model in representing the actual behavior. Simultaneously, the SEC stiffness ranged from 218 to 376 kN/m across the participants, which was slightly higher than the approximate 200 kN/m reported for the Achilles tendon (Ekiert et al., 2021; Kurihara et al., 2012; Lichtwark and Wilson, 2005). However, given the considerable variation in Achilles tendon stiffness owing to factors such as participants’ anatomy, level and type of physical activity (Lamontagne and Kennedy, 2013), and measurement methodologies (Finni and Vanwanseele, 2023), these differences remain within reasonable limits. These results regarding the SEC stiffness suggest that our MTC model captures the dynamics of the MTC of the MG muscle.

Interestingly, the CC exhibited a negative stiffness as the hop frequency increased (Fig. 6). These findings may appear strange because stiffness is generally defined for objects passively stretched under a force and has only positive values. However, the active force exerted by the muscles introduces a novel interpretation. According to the definition of stiffness, which is the ratio of force to elongation, muscle stiffness becomes negative when a muscle concentrically contracts against an increasing load. Using Eq. 1, we can derive the following expression for calculating MTC stiffness: *k*_*MTC*_ = *k*_*CC*_ ⋅ *k*_*SEC*_ /(*k*_*CC*_ + *k*_*SEC*_ ) . This equation indicates that MTC stiffness can exceed the SEC stiffness only for a negative CC stiffness (Fig. 7(A)). These results may appear to diverge from our hypothesis that a higher CC stiffness leads to a higher MTC stiffness. However, expressing MTC stiffness as a function of the inverse of CC stiffness, *k*_*MTC*_ = *k*_*SEC*_/(1 + *k*_*SEC*_ /*k*_*CC*_ ) , provides a unified interpretation of the negative CC stiffness as an extension of a very large positive stiffness (Fig. 7(B)). In summary, MTC stiffness equals SEC stiffness for an isometric CC, decreases for an elongating CC, and increases for a shortening CC. In this study, instructions to minimize the contact duration required the participants to maximize the MTC stiffness, thereby leading to the observation of a negative CC stiffness.

**Fig. 7.**
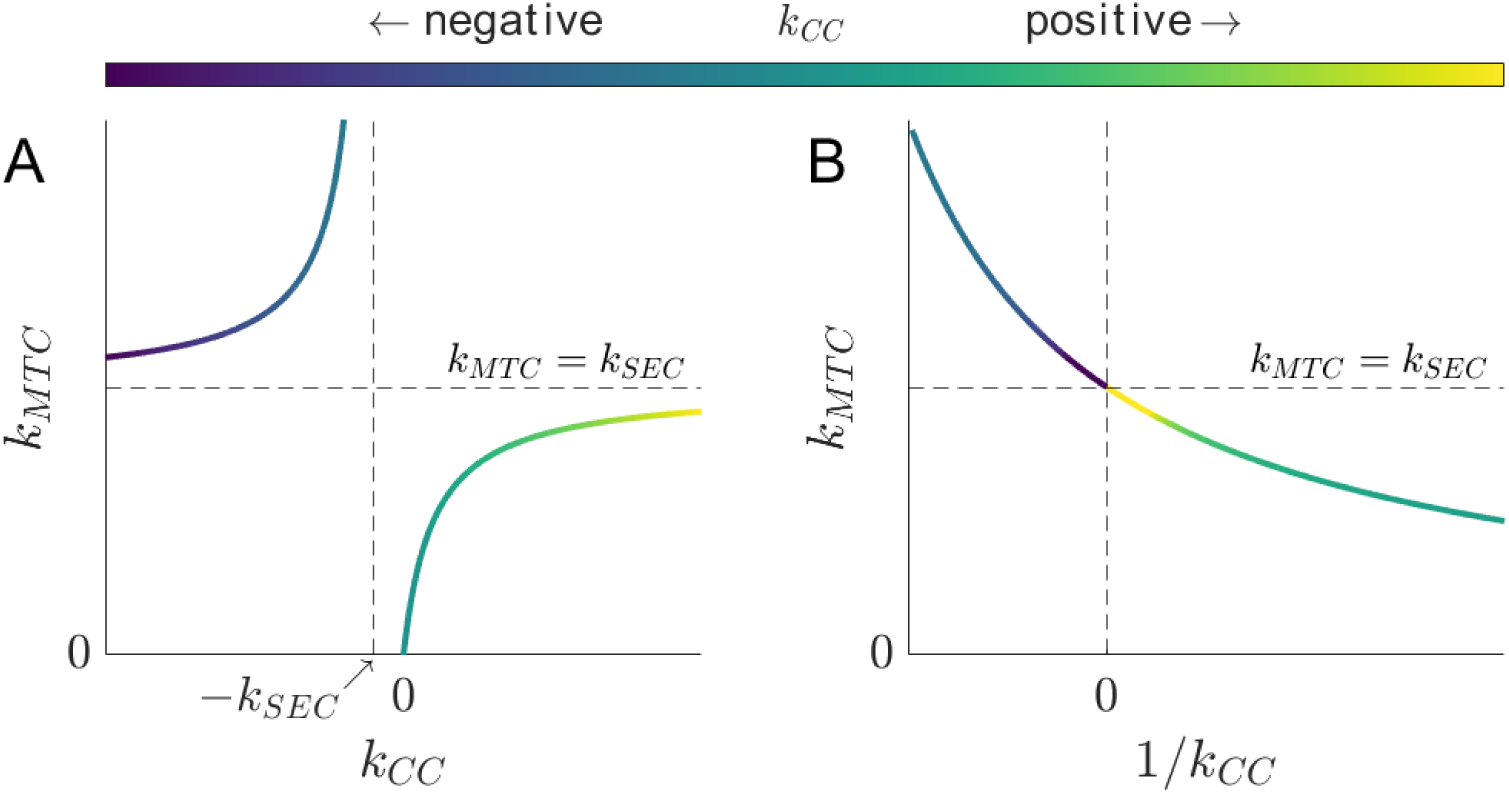
Relationship between MTC and CC stiffnesses based on Eq. 1. MTC stiffness is represented as a function of CC stiffness (A) and the inverse of CC stiffness (B). The colors of each curve represent CC stiffness values. Although MTC stiffness never exceeds SEC stiffness for a positive CC stiffness (the lower right part of each panel), it exceeds SEC stiffness for a negative CC stiffness (the upper left part of each panel).

Our model offers a unified framework for interpreting the results of previous studies that examined fascicle dynamics during hopping. For example, Jessup et al. (2023) showed that the lateral gastrocnemius muscle lengthens at low hop frequencies but remains almost isometric at high frequencies. Based on these results, CC stiffness appears to increase with hop frequency. Although their results differ from ours in that they revealed lengthening fascicles at low hop frequencies, which is attributed to differences in participant-related factors, their results align with ours in that the CC stiffness contributed to a higher MTC stiffness required at higher hop frequencies. Another study on hopping at different intensities (Hoffrén et al., 2012, 2011) showed that the MG shortens during hopping at higher intensities where a higher leg stiffness is required. Notably, their finding that muscle shortening during the braking phase of hopping is associated with increased ankle joint stiffness is equivalent to our concept of negative muscle stiffness.

Our explanation that the increased MTC stiffness is due to the fascicle shortening raises a question: why was the isometric contraction of the fascicles observed at a hop frequency of 2 Hz? The participant could have increased the MTC stiffness to reduce the contact duration by shortening the fascicle. The observed isometric behavior can be attributed to the load placed on the muscle. As shown in Fig. 4, the contact duration was relatively constant across different hop frequencies, whereas the aerial duration was longer at lower frequencies. To obtain a larger impulse required for a longer aerial duration while maintaining the same contact duration, the GRF must also be increased. Overall, the muscle was loaded more heavily at lower frequencies than at higher frequencies. Owing to the force–velocity relationship of the muscle (Hill, 1938), shortening against such a heavy load is unfavorable for the muscle to generate forces. This explains why the participants were unable to reduce further their contact duration by shortening the fascicle at 2 Hz. However, if the participants had been subjected to even lower frequencies with greater muscle loads, eccentric contractions might have been observed to endure the load. This prediction aligns with the results of a previous study that compared drop jumps at different intensities (Sousa et al., 2007). Thus, a frequency of approximately 2 Hz used in this study could represent the turning point where muscle dynamics change from concentric to eccentric contractions.

The isometric behavior of the fascicles has been implicated in movement with SSC as an energy-saving mechanism (Biewener et al., 1998; Fukunaga et al., 2001; Roberts et al., 1997). In cyclic calf-raise exercises, which are characterized by repetitive vertical movements of the CoM similar to hopping, such isometric behavior has been identified as resonance in the triceps surae MTC (Bach et al., 1983; Dean and Kuo, 2011; Takeshita et al., 2006). Resonance occurs when the movement matches the resonant frequency of the MTC, and the situation where the tendon becomes responsible for the change in the length of the entire MTC is equivalent to the oscillation of the mass-spring system, with the stiffness of its spring being similar to that of the SEC (Fig. 7).

Applying this concept to hopping suggests the existence of a “resonant hop frequency,” where fascicles remain isometric during ground contact. However, the MTC resonance cannot be uniquely determined solely by hop frequency because of the flexibility in selecting the contact and aerial durations for the same hop frequency. With the analogy of the cyclic calf raise, hopping with an isometric fascicle length should be equivalent to that of the mass-spring system, with the stiffness of its spring being the same as that of the SEC. The contact duration necessary to achieve this isometric condition is expected to decrease as the aerial duration increases (Blickhan, 1989). In fact, in this study, the contact duration during hopping at 2 Hz (181 ms), where isometric contraction of the muscle was observed, was considerably shorter than the contact duration corresponding to the resonant frequency of the previously reported calf raise exercise (300–333 ms). Therefore, we speculate that for a given hop frequency, isomeric contraction of the fascicles during ground contact can be achieved by selecting a proper combination of contact and aerial durations.

The MTC model-based framework presented in this study can be applied to other SSC exercises, such as walking and running. During walking and running, variations in fascicle behavior owing to movement speed have been reported (Farris and Sawicki, 2012; Ishikawa and Komi, 2007; Lai et al., 2014; Swinnen et al., 2018, 2022), which can be explained as in the present study. However, walking and running exhibit higher degrees of freedom than hopping and require modifications when the MTC model is applied to understand the behavior of the fascicles. First, although hopping was approximated as a vertical movement in this study, walking and running involve simultaneous horizontal movements. Second, because using a single MTC model to describe whole-body movement is inadequate for modeling multi-joint motion, a multi-body model must be constructed (Nikooyan and Zadpoor (2011)). Future studies are warranted to reveal the relationship between the stiffness of multiple muscles and movement parameters such as contact duration and locomotion speed.

This study has several limitations. First, knee stiffness may have partially contributed to leg stiffness. However, we believe that the effect of the knee was small under our experimental conditions because of the strong correlation observed between ankle and leg stiffnesses (with correlation coefficients exceeding 0.91 and *p*-values less than 0.01, except for one participant) as well as the lack of a statistically significant correlation between the knee and leg stiffnesses for all participants (with *p*-values above 0.098), as reported in a previous study (Hobara et al., 2011). Moreover, the smaller range of motion and moment arm of the knee compared with those of the ankle resulted in a smaller contribution to the MTC length (3.5 cm from the ankle and 0.7 cm from the knee at 2 Hz).

Another limitation is that the dynamics of the MG were studied as a representative of the triceps surae muscle without considering the lateral gastrocnemius or soleus muscles. Furthermore, the effect of the antagonistic tibial anterior muscle (TA) was ignored. Although direct evidence regarding the dynamics of individual muscles during hopping is lacking, insights from cyclic calf-raise exercises (Sakuma et al., 2012) suggest that the frequency-dependent behavior among the muscles may be similar during hopping. In addition, TA co-contraction plays a minor role in leg stiffness control during hopping (Hobara et al., 2007). Consequently, substituting the body behavior during hopping with MG dynamics can be justified to a certain extent.

In conclusion, this study provides a quantitative explanation of leg stiffness adjustment governed by lower limb muscles during hopping. In particular, the negative stiffness of the CC makes the MTC stiffer than the SEC, thereby achieving the high leg stiffness required for high hop frequencies. This insight into the dynamic interaction between the muscles and tendons holds promising implications beyond the scope of this study, particularly for bouncing movements involving the SSC, such as running.

## Acknowledgements

This work was supported by the International Graduate Program of Innovation for the Intelligent World of the University of Tokyo and by JSPS KAKENHI (Grant Numbers JP19K24281, JP20K11330, and JP23K10684). We would like to thank Editage (www.editage.com) for English language editing. We would also like to thank the members of the Sports Biomechanics Lab at the University of Tokyo for their support during the experiments and valuable discussions.

## Declaration of generative AI and AI-assisted technologies in the writing process

During the preparation of this work, the authors used ChatGPT in order to enhance the quality of language and readability. After using this tool/service, the authors reviewed and edited the content as needed and take full responsibility for the content of the publication.

## CrediT Author contribution statement

**Kazuki Kuriyama**: Conceptualization, Data curation, Formal Analysis, Funding acquisition, Investigation, Methodology, Project administration, Resources, Software, Validation, Visualization, Writing – original draft, Writing – review & editing. **Daisuke Takeshita**: Conceptualization, Funding acquisition, Project administration, Supervision, Validation, Writing - review & editing.

## Data availability statement

The datasets underlying the current study are available on request.

## References

Ae, M., Tang, H., Yokoi, T., 1992. Estimation of inertia properties of the body segments in Japanese athletes. Biomechanisms 11, 23–33.

Arampatzis, A., Brüggemann, G.P., Metzler, V., 1999. The effect of speed on leg stiffness and joint kinetics in human running. Journal of Biomechanics 32, 1349–1353.

Bach, T.M., Chapman, A.E., Calvert, T.W., 1983. Mechanical resonance of the human body during voluntary oscillations about the ankle joint. Journal of Biomechanics 16, 85–90.

Biewener, A.A., Konieczynski, D.D., Baudinette, R. V, 1998. In vivo muscle force-length behavior during steady-speed hopping in tammar wallabies. The Journal of experimental biology 201, 1681–94.

Blickhan, R., 1989. The spring-mass model for running and hopping. Journal of Biomechanics 22, 1217–1227.

Bobbert, M.F., Huijing, P.A., van Ingen Schenau, G.J., 1986. A model of the human triceps surae muscle-tendon complex applied to jumping. Journal of Biomechanics 19, 887–898.

Brughelli, M., Cronin, J., 2008. Influence of Running Velocity on Vertical, Leg and Joint Stiffness. Sports Medicine 38, 647–657.

Butler, R.J., Crowell, H.P., Davis, I.M.C., 2003. Lower extremity stiffness: Implications for performance and injury. Clinical Biomechanics 18, 511–517.

Dean, J.C., Kuo, A.D., 2011. Energetic costs of producing muscle work and force in a cyclical human bouncing task. Journal of Applied Physiology 110, 873–880.

Ekiert, M., Tomaszewski, K.A., Mlyniec, A., 2021. The differences in viscoelastic properties of subtendons result from the anatomical tripartite structure of human Achilles tendon - ex vivo experimental study and modeling. Acta Biomaterialia 125, 138–153.

Farley, C.T., Blickhan, R., Saito, J., Taylor, C.R., 1991. Hopping frequency in humans: A test of how springs set stride frequency in bouncing gaits. Journal of Applied Physiology 71, 2127–2132.

Farley, C.T., Morgenroth, D.C., 1999. Leg stiffness primarily depends on ankle stiffness during human hopping. Journal of Biomechanics 32, 267–273.

Farris, D.J., Sawicki, G.S., 2012. Human medial gastrocnemius force-velocity behavior shifts with locomotion speed and gait. Proceedings of the National Academy of Sciences of the United States of America 109, 977–982.

Ferris, D.P., Farley, C.T., 1997. Interaction of leg stiffness and surface stiffness during human hopping. Journal of Applied Physiology 82, 15–22.

Ferris, D.P., Louie, M., Farley, C.T., 1998. Running in the real world: Adjusting leg stiffness for different surfaces. Proceedings of the Royal Society B: Biological Sciences 265, 989–994.

Finni, T., Vanwanseele, B., 2023. Towards modern understanding of the Achilles tendon properties in human movement research. Journal of Biomechanics 152, 111583.

Fukashiro, S., Noda, M., Shibayama, A., 2001. In vivo determination of muscle viscoelasticity in the human leg. Acta Physiologica Scandinavica 172, 241–248.

Fukunaga, T., Kubo, K., Kawakami, Y., Fukashiro, S., Kanehisa, H., Maganaris, C.N., 2001. In vivo behaviour of human muscle tendon during walking. Proceedings of the Royal Society B: Biological Sciences 268, 229–233.

Garciá-Pinillos, F., Garciá-Ramos, A., Ramírez-Campillo, R., Latorre-Román, P., Roche-Seruendo, L.E., 2019. How Do Spatiotemporal Parameters and Lower-Body Stiffness Change with Increased Running Velocity? A Comparison between Novice and Elite Level Runners. Journal of Human Kinetics 70, 25–38.

Hawkins, D., Hull, M.L., 1990. A method for determining lower extremity muscle-tendon lengths during flexion/extension movements. Journal of Biomechanics 23, 487–494.

Hill, A.V., 1938. The heat of shortening and the dynamic constants of muscle. Proceedings of the Royal Society of London. Series B - Biological Sciences 126, 136–195.

Hobara, H., Inoue, K., Omuro, K., Muraoka, T., Kanosue, K., 2011. Determinant of leg stiffness during hopping is frequency-dependent. European Journal of Applied Physiology 111, 2195–2201.

Hobara, H., Kanosue, K., Suzuki, S., 2007. Changes in muscle activity with increase in leg stiffness during hopping. Neuroscience Letters 418, 55–59.

Hoffrén, M., Ishikawa, M., Avela, J., Komi, P. V., 2012. Age-related fascicle-tendon interaction in repetitive hopping. European Journal of Applied Physiology 112, 4035–4043.

Hoffrén, M., Ishikawa, M., Rantalainen, T., Avela, J., Komi, P. V., 2015. Neuromuscular mechanics and hopping training in elderly. European Journal of Applied Physiology 115, 863–877.

Hoffrén, M., Ishikawa, M., Rantalainen, T., Avela, J., Komi, P. V., 2011. Age-related muscle activation profiles and joint stiffness regulation in repetitive hopping. Journal of Electromyography and Kinesiology 21, 483–491.

Ishikawa, M., Finni, T., Komi, P. V., 2003. Behaviour of vastus lateralis muscle-tendon during high intensity SSC exercises in vivo. Acta Physiologica Scandinavica 178, 205–213.

Ishikawa, M., Komi, P. V., 2007. The role of the stretch reflex in the gastrocnemius muscle during human locomotion at various speeds. Journal of Applied Physiology 103, 1030–1036.

Jessup, L.N., Kelly, L.A., Cresswell, A.G., Lichtwark, G.A., 2023. Linking muscle mechanics to the metabolic cost of human hopping. Journal of Experimental Biology 226.

Kubo, K., Kawakami, Y., Fukunaga, T., 1999. Influence of elastic properties of tendon structures on jump performance in humans. Journal of Applied Physiology 87, 2090–2096.

Kuitunen, S., Ogiso, K., Komi, P. V., 2011. Leg and joint stiffness in human hopping. Scandinavian Journal of Medicine & Science in Sports 21.

Kurihara, T., Sasaki, R., Isaka, T., 2012. Mechanical properties of the Achilles tendon in relation to various sport activities of collegiate athletes, In Proceedings of the 30th International Conference on Biomechanics in Sports. Melbourne, Australia. 2012.

Lai, A., Schache, A.G., Lin, Y.C., Pandy, M.G., 2014. Tendon elastic strain energy in the human ankle plantar-flexors and its role with increased running speed. Journal of Experimental Biology 217, 3159–3168.

Lamontagne, M., Kennedy, M.J., 2013. The biomechanics of vertical hopping: A review. Research in Sports Medicine 21, 380–394.

Lichtwark, G.A., Wilson, A.M., 2005. In vivo mechanical properties of the human Achilles tendon during one-legged hopping. Journal of Experimental Biology 208, 4715–4725.

McMahon, T.A., Cheng, G.C., 1990. The mechanics of running: How does stiffness couple with speed? Journal of Biomechanics 23, 65–78.

Monte, A., Nardello, F., Zamparo, P., 2021. Mechanical advantage and joint function of the lower limb during hopping at different frequencies. Journal of Biomechanics 118, 110294.

Morin, J.B., Jeannin, T., Chevallier, B., Belli, A., 2006. Spring-mass model characteristics during sprint running: Correlation with performance and fatigue-induced changes. International Journal of Sports Medicine 27, 158–165.

Nikooyan, A.A., Zadpoor, A.A., 2011. Mass–spring–damper modelling of the human body to study running and hopping – an overview. Proceedings of the Institution of Mechanical Engineers, Part H: Journal of Engineering in Medicine 225, 1121–1135.

Roberts, T.J., Marsh, R.L., Weyand, P.G., Taylor, C.R., 1997. Muscular force in running turkeys: The economy of minimizing work. Science 275, 1113–1115.

Sakanaka, T.E., Gill, J., Lakie, M.D., Reynolds, R.F., 2018. Intrinsic ankle stiffness during standing increases with ankle torque and passive stretch of the Achilles tendon. PLoS ONE 13.

Sakuma, J., Kanehisa, H., Yanai, T., Fukunaga, T., Kawakami, Y., Moritani, T., 2012. Fascicle-tendon behavior of the gastrocnemius and soleus muscles during ankle bending exercise at different movement frequencies. European Journal of Applied Physiology 112, 887–898.

Sano, K., Ishikawa, M., Nobue, A., Danno, Y., Akiyama, M., Oda, T., Ito, A., Hoffrén-Mikkola, M., Nicol, C., Locatelli, E., Komi, P. V., 2013. Muscle-tendon interaction and EMG profiles of world class endurance runners during hopping. European Journal of Applied Physiology 113, 1395–1403.

Shorten, M.R., 1987. Muscle Elasticity and Human Performance, Current Reserch in Sports Biomechanics. Karger Publishers, pp. 1–18.

Sousa, F., Ishikawa, M., Vilas-Boas, J.P., Komi, P. V., 2007. Intensity- and muscle-specific fascicle behavior during human drop jumps. Journal of Applied Physiology 102, 382–389.

Struzik, A., Karamanidis, K., Lorimer, A., Keogh, J.W.L., Gajewski, J., 2021. Application of Leg, Vertical, and Joint Stiffness in Running Performance: A Literature Overview. Applied Bionics and Biomechanics.

Swinnen, W., Hoogkamer, W., Delabastita, T., Aeles, J., De Groote, F., Vanwanseele, B., 2018. Effect of habitual foot-strike pattern on the gastrocnemius medialis muscle-tendon interaction and muscle force production during running. Journal of Applied Physiology 126, 708–716.

Swinnen, W., Mylle, I., Hoogkamer, W., De Groote, F., Vanwanseele, B., 2022. Triceps surae muscle force potential and force demand shift with altering stride frequency in running. Scandinavian Journal of Medicine and Science in Sports 32, 1444–1455.

Takeshita, D., Shibayama, A., Muraoka, T., Muramatsu, T., Nagano, A., Fukunaga, T., Fukashiro, S., 2006. Resonance in the human medial gastrocnemius muscle during cyclic ankle bending exercise. Journal of Applied Physiology 101, 111–118.

Van Hooren, B., Bosch, F., 2016. Influence of Muscle Slack on High-Intensity Sport Performance: A Review. Strength & Conditioning Journal 38, 75–87.

Van Hooren, B., Teratsias, P., Hodson-Tole, E.F., 2020. Ultrasound imaging to assess skeletal muscle architecture during movements: a systematic review of methods, reliability, and challenges. Journal of Applied Physiology 128, 978–999.

Wakeling, J.M., Febrer-Nafría, M., De Groote, F., 2023. A review of the efforts to develop muscle and musculoskeletal models for biomechanics in the last 50 years. Journal of Biomechanics.

Winter, D.A., 2009. Biomechanics and Motor Control of Human Movement. John Wiley & Sons, Inc., Hoboken, NJ, USA.

